# Predicting establishment risk of the Korean perch *Coreoperca herzi* in Japan using species distribution modelling: Insights for invasive fish management

**DOI:** 10.64898/2026.02.21.707239

**Authors:** Subaru Morimoto, Satsuki Tsuji

## Abstract

Invasive freshwater fishes pose serious threats to native biodiversity and ecosystem functioning, but management resources for prevention and control are limited. The Korean perch, *Coreoperca herzi*, a predatory freshwater fish native to the Korean Peninsula, has recently established itself in the Oyodo and Tone River system in Japan and has begun causing serious negative impacts on the ecosystem. This study aimed to predict the spatial distribution of potential establishment risk of the Korean perch within the Japanese archipelago by using a species distribution model based on occurrence records from the native range and environmental variables. Presence records were obtained from the Global Biodiversity Information Facility and combined with hydroclimatic variables from WorldClim and HydroATLAS. A random forest algorithm modified for class-imbalanced data was employed. Model tuning and evaluation were conducted using catchment-based spatial cross-validation to enhance spatial transferability, and predictive performance was assessed using the area under the curve (AUC) and the true skill statistic (TSS). The final model achieved high predictive accuracy (AUC = 0.80; TSS = 0.56) on an independent test dataset. River discharge, river slope, and maximum summer water temperature were identified as key predictors of habitat suitability. Projection of the model onto Japanese river networks indicated that environmentally suitable habitats for Korean perch are widely distributed, particularly in mid- to upstream reaches of rivers. These results provide a spatial basis for prioritising monitoring and eradication and highlight the importance of preventive management in high-risk catchments before invasion or at early invasion stages.

## Introduction

Invasive species pose a serious threat to native ecosystems, economic activities such as agriculture and fisheries, and the health of both humans and wildlife (Diagne et al., 2021; Early et al., 2016; Gallardo et al., 2016; Turbelin et al., 2024). The intensification of global trade, transportation networks, and human mobility has markedly increased both the frequency and geographic scale of biological invasions (Dukes & Mooney, 1999; Gippet et al., 2019; Seebens et al., 2021). As a result, preventing new introductions and mitigating their impacts have become urgent priorities in conservation science and environmental policy (Simberloff et al., 2013). Freshwater ecosystems, in particular, are relatively closed and are also deeply linked to human activities (Faghihinia et al., 2021; Sato et al., 2010). Consequently, they face a high risk of introduction and establishment of invasive species and are vulnerable to the threats they pose (Cox & Lima, 2006; Havel et al., 2015). However, the presence of aquatic organisms is often difficult to detect, and anthropogenic releases are largely unpredictable (Havel et al., 2015). Under such constraints, comprehensive surveillance across all water bodies is rarely feasible. Therefore, spatially explicit risk assessment that identifies areas with high establishment probability is essential for prioritising monitoring and management efforts under limited budgets and personnel (McGeoch et al., 2016a).

The Korean perch (*Coreoperca herzi* Herzenstein, 1896; Fig. 1a) is a freshwater fish native to the Korean Peninsula that exemplifies these challenges. The species was first recorded in Japan in 2017 in the Hagiwara River, a tributary of the Oyodo River system (Byeon, 2017; Hibino et al., 2019; Fig. 3b). Since then, it has shown a high predatory nature, invasiveness, and dispersal ability, steadily increasing both its biomass and distribution range within the river system (Hibino et al., 2022; Tsuji et al., 2024).

**Fig. 1.**
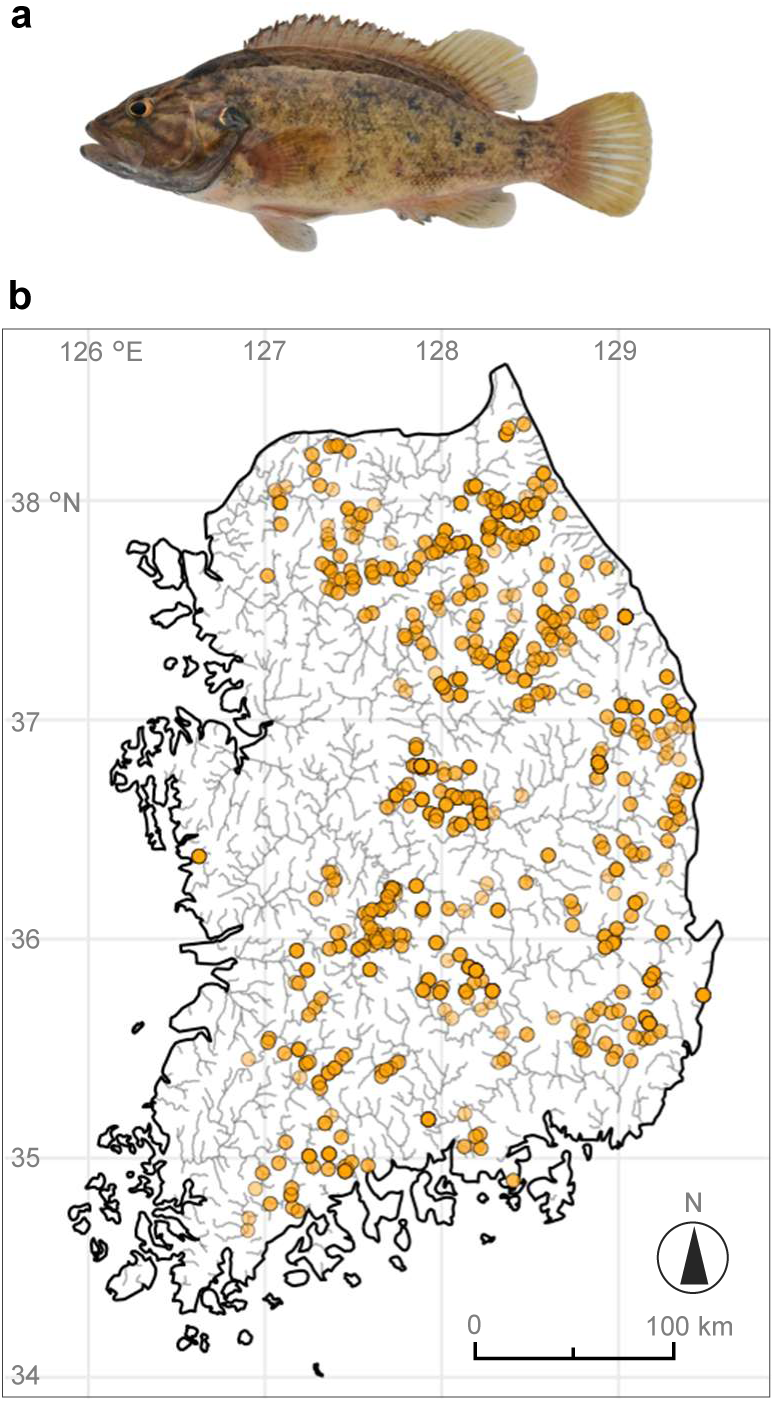
(a) Korean perch (*Coreoperca herzi* Herzenstein, 1896). (b) Locations of occurrence records for the Korean perch in South Korea from 2010 to 2024, obtained from the Global Biodiversity Information Facility (GBIF). A total of 1,221 locations existing on the river network obtained from HydroATLAS are shown.

This species actively preys on fish and aquatic insects and prefers mid- to upstream habitats with clean water and structurally complex refuges (Chae et al., 2019). It also has ecological traits that facilitate establishment, such as territorial behaviour and egg guarding during the breeding season (Chae et al., 2019; Uchida, 1935). These traits can enhance reproductive success and recruitment while intensifying predation pressure on native species. Indeed, the species has become dominant in parts of the Hagiwara River and has been associated with significant declines in native benthic fish populations (Hibino et al., 2022; Tsuji et al., 2024). The Korean perch can attain a total length of approximately 30 cm (Hibino et al., 2022; Uchida, 1935), enabling it to prey on a broad size range of native species. Furthermore, Japanese freshwater systems lack persistent top predators or strong competitors that could effectively regulate perch population growth (Hosoya, 2019), potentially facilitating population expansion and trophic imbalances. Therefore, the Korean perch is regarded with concern as having the potential to cause serious damage to freshwater ecosystems across Japan through continued range expansion.

Despite these concerns, the Korean perch is popular for ornamental purposes and recreational angling, which greatly increases the risk of human-mediated new and secondary translocation. Indeed, within the Oyodo River system, range expansion has been recognised at multiple sites beyond large weirs that lack fishways, which are impassable to natural dispersal (Tsuji et al., 2024). Furthermore, a single individual was first detected in 2024 in the Ayu River, a tributary of the Tone River system located several hundred kilometres from the Oyodo River system (Ministry of the Environment, 2025; Fig. 3b), and 101 individuals were subsequently captured at the same site in 2025, indicating rapid population establishment and expansion. Such occurrences that cannot be explained by natural dispersal strongly suggest anthropogenic introduction, raising concerns about the existence of unconfirmed populations and further spread. In response to the escalating situation, although local and national authorities are advancing regulations to prohibit transport, possession, and release of the Korean perch, completely preventing illegal or unintended releases is likely to be extremely difficult (McEachran et al., 2022). Under these circumstances, it would be advisable for management strategies to extend beyond prevention alone and incorporate proactive, risk-based approaches grounded in the precautionary principle.

For species with a higher invasion risk, such as the Korean perch, developing prioritised monitoring and eradication strategies that prioritise areas with high establishment risk is effective for achieving efficient control with limited budgets, personnel, and effort (McGeoch et al., 2016a). One of the most promising tools for this purpose is the species distribution model (SDM) (Rodríguez-Castañeda et al., 2012). SDMs predict a species’ potential distribution range based on the relationship between known occurrence records and environmental variables (Guisan & Thuiller, 2005; Guisan & Zimmermann, 2000). SDMs have been demonstrated to have high accuracy and practical utility in assessing the risks of invasion and establishment by non-native species (Elith et al., 2010a; Thuiller et al., 2005). In particular, the adoption of machine learning-based approaches such as random forest (RF) and maximum entropy (MaxEnt) has enabled the incorporation of complex environmental responses into predictions that were difficult to represent using conventional approaches (Elith et al., 2006; Olden et al., 2008; Valavi et al., 2021). Projecting SDMs across regions assumes ecological niche conservatism. Because parts of the Korean Peninsula and Japan share similar climatic and hydrological conditions, native-range data should offer a reasonable basis for estimating establishment risk in Japan, albeit with inherent transferability uncertainty.

This study aims to predict the spatial distribution of potential establishment risk of the Korean perch within Japan by constructing an SDM based on occurrence records and environmental variables from South Korea and projecting it onto the Japanese archipelago. By clarifying areas with a high risk of establishment, the results of this study will enable the prioritisation of monitoring and eradication efforts based on scientific evidence. Ultimately, these findings are expected to contribute to the development of strategic management policies aimed at preventing the future spread of the Korean perch and conserving native freshwater ecosystems.

## Materials and Methods

### Acquisition of occurrence records and environmental variables

Occurrence records of the Korean perch, recorded between 2010 and 2024, were obtained from the Global Biodiversity Information Facility (GBIF) for 1,221 locations within South Korea (Fig. 1b). It is recognised that large open-access databases such as GBIF inherently contain spatial sampling biases, including repeated surveys at the same locations and preferential sampling of easily accessible areas. To reduce such spatial biases, spatial thinning was employed, and duplicate occurrence records located within 5 km of each other were excluded. Pseudo-absence locations were randomly sampled from areas located at least 5 km away from occurrence records. Environmental variables were obtained from WorldClim ver. 2 (Fick & Hijmans, 2017) and HydroATLAS (Linke et al., 2019). All layers were unified to a common spatial resolution of 15 arc-seconds (approximately 500 m), using bilinear interpolation. The temperature data from WorldClim were converted to water temperature using the conversion formula by Punzet et al. (2012):

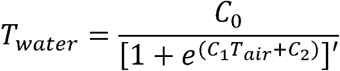

where *T*_water_ = water temperature (°C), *T*_air_ = air temperature (°C), *C*_0_ = upper bound water temperature (°C), *C*_1_ = steepest slope of the function (°C−1), and *C*_2_ = measure for inflexion point of the function (°C), (inflection point = −*C*_2_/*C*_1_). The coefficients of an s-shaped function have physical properties relating to the water–air temperature relationship. Elevation was excluded from the analysis because it can generally be regarded as an indirect proxy for habitat conditions, mediated through factors such as water temperature and discharge, rather than a direct determinant of habitat suitability for freshwater fishes. To minimise the effects of multicollinearity and ensure spatial transferability of the model, one variable from each pair with a Pearson correlation coefficient exceeding 0.75 was excluded (Fig. S1). Additionally, variables with a Variance Inflation Factor (VIF) greater than 10 were not adopted. Based on these criteria, together with considerations of the ecological plausibility of the Korean perch, five environmental variables were finally selected (Table S1): river discharge (dis_m3_pyr), the maximum water temperature of the warmest month (water_bio5), river slope (sgr_dk_rav), groundwater depth (gwt_cm_cav), and sand content in surface soil (snd_pc_cav). The analysis area was restricted to locations on river networks using the layer provided by HydroATLAS.

### Model construction

In this study, a species distribution model (SDM) of the Korean perch was constructed using the machine learning algorithm random forest (RF), based on occurrence records from their native range in South Korea. RF is well-suited to capturing nonlinear relationships and higher-order interactions among predictor variables (Cutler et al., 2007). Comprehensive benchmark studies of SDMs have reported that RF methods designed to address class imbalance, such as down-sampled RF, exhibit high predictive accuracy alongside Boosted Regression Trees (BRT) and MaxEnt (Valavi et al., 2022). However, RF implemented with default settings is vulnerable to severe class imbalance between presence and background data, as well as to overlap between these classes in environmental space (Valavi et al., 2021).

These issues can lead to overfitting, particularly through excessive adaptation to background data (Valavi et al., 2023). To address these limitations and improve model generalisability and spatial transferability, our RF model was modified following the recommendations of Valavi et al. (2021). First, during the construction of each decision tree, background data were randomly down-sampled to match the number of occurrence records, thereby suppressing biased partitioning caused by class imbalance. Second, the Hellinger distance was adopted as the node splitting criterion, which is more robust against imbalanced datasets, enabling appropriate partitioning even in environmental spaces with substantial class overlap. Third, to mitigate overfitting, the maximum depth of individual trees was constrained. Using shallow trees is expected to limit overly complex decision boundaries and enhance model generalisation. Finally, to ensure prediction stability, the number of trees was set to 2,000.

Model construction was conducted using R software (version 4.4.2; R Core Team, 2021) with the tidymodels framework and the tidysdm package (Leonardi et al., 2024). The RF algorithm was implemented using the ranger package (Wright & Ziegler, 2017), and probability forests were employed to obtain probabilistic predictions. The Hellinger distance was used as the node-splitting criterion, the maximum tree depth was set to 5, and the total number of trees was fixed at 2,000. Two hyperparameters were explored via Bayesian optimisation: mtry (the number of candidate variables considered at each node) and min_n (the minimum number of samples required for a split). As the evaluation metric, the area under the curve (AUC) was employed from internal cross-validation. AUC values range from 0 to 1, with a value of 0.5 indicating performance equivalent to random prediction. The optimisation process consisted of an initial evaluation of ten parameter sets, followed by up to fifty additional iterations. The exploration was terminated if no improvement in performance was observed over ten consecutive iterations.

### Catchment-based spatial cross-validation

To maximise model generalisability and spatial transferability, a nested spatial cross-validation scheme was implemented using catchment boundaries as the spatial partitioning units. Catchment delineations were derived from HydroBASINS (Lehner & Grill, 2013; https://www.hydrosheds.org/products/hydrobasins), which provides a hierarchical system of watershed divisions. In this study, level 8 basins were selected as an appropriate spatial scale that ensured sufficient spatial independence among folds while retaining an adequate number of occurrence records for stable model training and evaluation within each partition (Fig. S2a). First, the full dataset was partitioned by catchment into training (80%) and testing (20%) subsets. The testing dataset was withheld entirely from model fitting and hyperparameter tuning and was used exclusively for the final evaluation of model generalisability and spatial transferability. Next, the training dataset was further subjected to catchment-based grouped five-fold cross-validation (group k-fold CV, k = 5) for hyperparameter tuning (Fig. S2b). During model training, down-sampling was applied to address class imbalance between presence and background data, with background samples adjusted to match the number of presence records. Furthermore, to mitigate the effects of randomness associated with data partitioning and sampling, the entire workflow from splitting to model evaluation was repeated ten times.

### Model Evaluation

To evaluate the predictive performance of the constructed RF-based model, assessments were conducted based on variable importance, partial dependence plots (PDPs), and classification performance metrics. The relative importance of each explanatory variable was evaluated using permutation importance. Specifically, the decrease in AUC resulting from random permutation of the values of each variable was calculated, with larger decreases interpreted as indicating a greater contribution to model predictions. Variable importance was computed across multiple iterations of training, and the mean values were used for evaluation. In addition, PDPs were generated for each variable to visualise the relationship between each variable and the predicted habitat suitability potential under conditions where the influence of other variables was averaged out. Model classification performance was evaluated using AUC and True Skill Statistic (TSS). The model evaluations were performed using an independent test dataset not used for training or hyperparameter tuning, verifying generalisation performance. The TSS was calculated based on the optimal threshold maximising the sum of sensitivity and specificity. Finally, the spatial validity of model predictions was confirmed by comparing the spatial patterns of predicted habitat suitability potential with the distribution of known occurrence records within the native range in South Korea.

### Projection of the model onto the Japanese Archipelago

For each repetition, the RF model exhibiting the highest predictive performance was defined as the best RF model. The final ensemble prediction was then generated by calculating the arithmetic mean of predictions from these best-performing models. Before projecting the model onto the Japanese archipelago, a multivariate environmental similarity surface (MESS) analysis (Elith et al., 2010b) was conducted to reduce uncertainty associated with extrapolation beyond the range of environmental conditions represented in the training data. For each prediction location in the Japanese archipelago, environmental similarity was calculated relative to the training data derived from South Korea. Locations with negative MESS values (< 0) indicate novel environmental conditions not present in the training data. These locations were excluded from further analysis, and the habitat suitability potential of the Korean perch was evaluated only within the environmental domain in which the model was considered applicable.

## Results

Evaluation of variable importance using permutation importance indicated that river discharge exhibited the largest drop in AUC and was the most influential predictor in the model (Fig. 2a). Next, the maximum water temperature of the warmest month and river slope were found to have relatively high importance, and these variables also contributed significantly to habitat suitability prediction. In contrast, the importance of groundwater depth and sand content in surface soil was relatively small. PDPs for each environmental variable confirmed non-linear responses of predicted occurrence probability (Fig. 2b). For river discharge, habitat suitability was highest at low to medium flow levels, peaking at approximately 40 m³ s ¹, and gradually declined with increasing discharge. For the maximum water temperature of the warmest month, habitat suitability reached a maximum at around 25–26 °C, with a tendency to decrease at higher temperatures. Regarding river slope, habitat suitability was low in low-gradient reaches and relatively higher in reaches with moderate to high gradients.

**Fig. 2.**
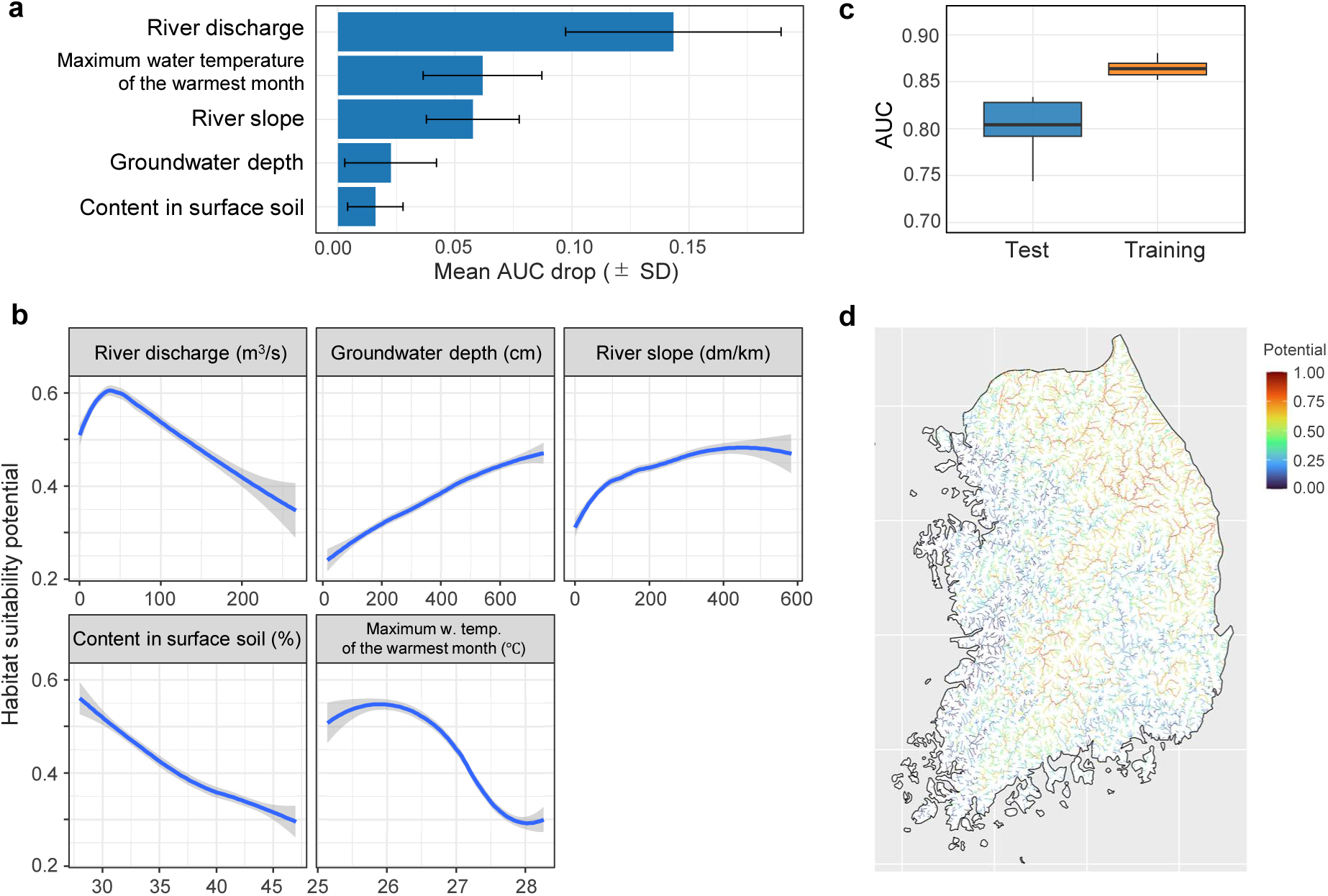
Summary of model performance evaluation results. (a) Evaluation results of environmental variable importance by AUC. The bar chart indicates the decrease in AUC (AUC drop; average ± SD) when each explanatory variable is randomly replaced. (b) Partial Dependence Plots (PDPs) for variables used in the model. Each curve represents the average PDP obtained from the model constructed at each iteration, smoothed using LOESS regression (with 95% confidence intervals). The vertical axis represents the model’s predicted value (habitat suitability potential) under conditions where the effects of other explanatory variables are averaged out. (c) AUC for the best model in each iteration on the training dataset (Training; orange) and the independent test dataset (Test; blue). (d) Results of projecting the predicted habitat suitability potential onto South Korea, the native range of the Korean perch, using the constructed SDM. Colour indicates habitat suitability potential (0–1), with red denoting higher suitability

The constructed RF model achieved an AUC of 0.86 ± 0.01 (mean ± SD) on the training dataset and an AUC of 0.80 ± 0.03 on the independent test dataset, which was not used for model training or hyperparameter tuning (Fig. 2c). Additionally, the threshold-dependent evaluation metric, the true skill statistic (TSS), was 0.56 ± 0.02. The optimal threshold used to calculate TSS was 0.51 ± 0.03. Estimating habitat suitability using the constructed model for the native range of the Korean perch in South Korea showed relatively high suitability in the central and western mid-to-upper reaches, where numerous occurrence records have been reported (Fig. 2d).

MESS analysis was used to assess the similarity of environmental conditions between South Korea (the native range of the Korean perch) and Japan, and positive MESS values were observed across most of Honshu, Shikoku, and Kyushu, indicating that environmental conditions in these areas fell within the range represented by the training data (Fig. 3a). In contrast, negative MESS values were detected in parts of north-eastern Hokkaido, some lowland areas of Honshu, and portions of the southwestern islands, indicating the presence of environmental conditions not represented in the native distribution of the Korean perch in South Korea. Overall, a large proportion of riverine areas in the Japanese archipelago exhibited similar environmental conditions to those in South Korea.

**Fig.3.**
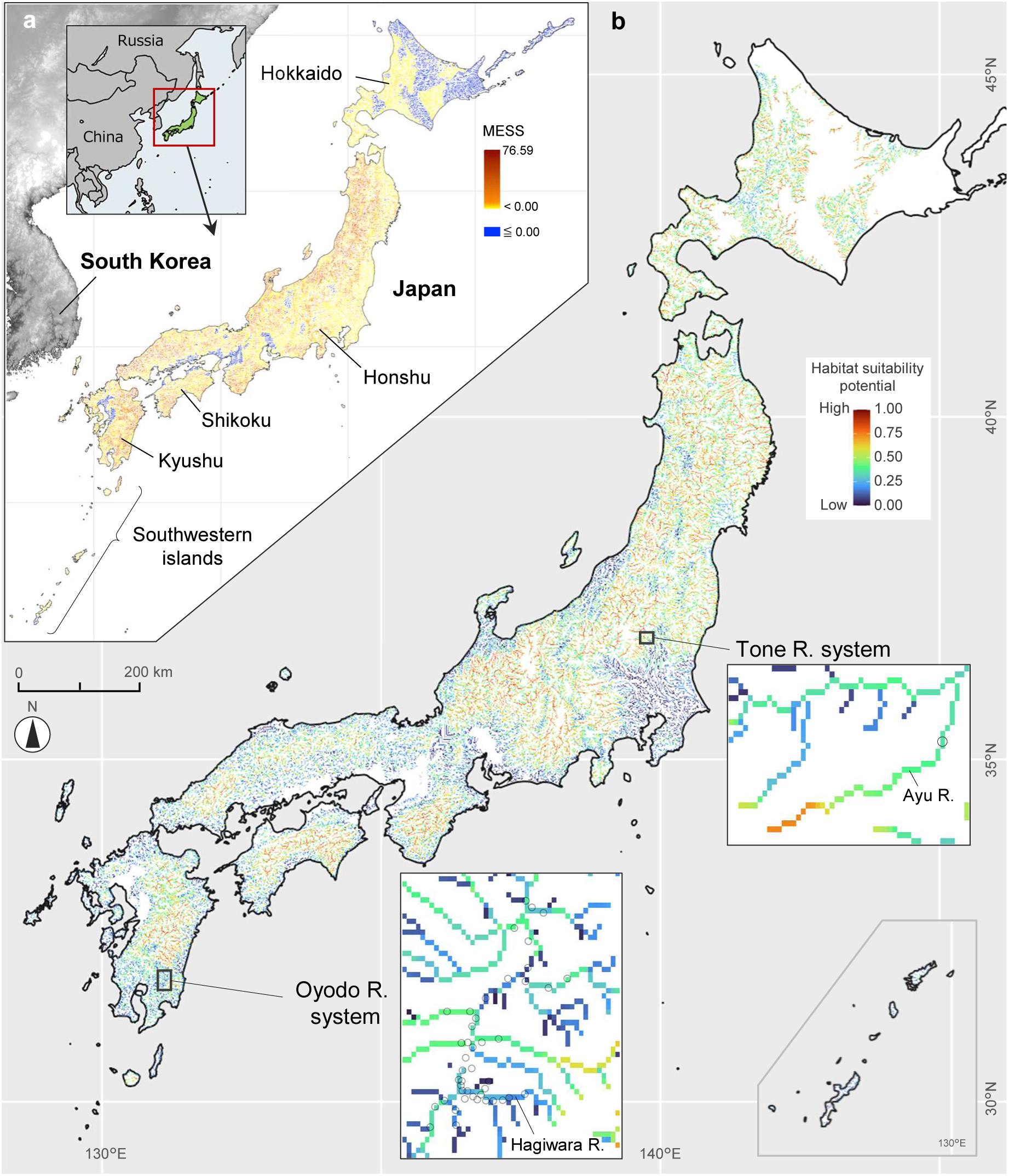
(a) Multivariate environmental similarity surface (MESS) score map for the Japanese Archipelago. Positive values indicate conditions similar to the training data in South Korea, while negative values indicate conditions outside the range of the training data. (b) Habitat suitability potential for the Korean perch in the Japanese archipelago, estimated using the constructed SDM. Areas encompassing the upper reaches of the Oyodo River system and the Ayu River in the Tone River system are enlarged, with locations where the Korean perch has been detected in previous studies indicated by circles

Projection of the constructed RF model onto the Japanese archipelago showed that areas with high habitat suitability potential for the Korean perch were widely distributed (Fig. 3b). In particular, river sections in inland and mountainous regions of Honshu, Shikoku, and Kyushu exhibited moderate to high suitability. On the other hand, relatively low suitability was predicted for many coastal and lowland areas. In the upper reaches of the Oyodo River system, which represents an established invasion area of the Korean perch, predicted habitat suitability was generally low to moderate (approximately 0.2 to 0.6). In the Ayu River within the Tone River system, where new invasions were confirmed in 2024, the predicted habitat suitability was moderate to high (approximately 0.35 to 0.75) with an increase in potential observed further upstream.

## Discussion

### Spatial distribution of habitat suitability potential for the Korean perch

The SDM developed in this study suggests that environmentally suitable conditions for the Korean perch are present across mid- to upper reaches of river systems within the Japanese archipelago. Given the model’s predictive performance (AUC and TSS values indicating acceptable discrimination ability), these projections provide a spatially explicit estimate of potential establishment risk. However, predictions should be interpreted as relative suitability rather than definitive evidence of future invasion. Within this context, the mapped distribution of suitable habitat can be regarded as an early warning indicator to support precautionary management.

River discharge, river slope, and summer water temperature were identified as the primary environmental factors governing the distribution of suitable habitat for the Korean perch. These variables are ecologically interpretable in light of the species’ life history traits and behavioural characteristics. In mid- to upper-reaches characterised by relatively low discharge and moderate slope, riverbeds often consist of gravel- and cobble-dominated substrates that promote structurally complex benthic habitats (Church, 2002). Such environments are likely to provide favourable conditions for the Korean perch, which exhibits territorial behaviour and employs an ambush-based predation strategy (Chae et al., 2019). Additionally, stable flow conditions and the availability of refuges are important determinants of reproductive success for this species, in which males construct nests and guard eggs during the breeding season (Chae et al., 2019; Uchida, 1935). These habitat characteristics are consistent with those observed in their native range on the Korean Peninsula, supporting the ecological plausibility of the environmental predictors identified for the model. Furthermore, the response patterns to water temperature should also be noted.

Although the Korean perch is not a cold-water species, it is also not adapted to extremely high water temperatures, and our predictions suggested that optimal conditions occur at maximum temperatures of the warmest month, around 25–26 °C. This temperature range is typical of mid-mountain rivers with abundant vegetation and shading (Poole & Berman, 2001). In natural river systems, discharge, slope, and thermal regimes often co-vary spatially, potentially generating overlapping environmental conditions favourable to the Korean perch. This interpretation refers to the ecological co-occurrence of environmental gradients rather than statistically tested interaction effects within the model.

The spatial distribution of habitat suitability potential revealed by the SDM, together with the ecological interpretation of the underlying environmental drivers, is expected to provide insights applicable to the design of control strategies and management practices at the field level. At present, confirmed occurrences of the Korean perch in Japan remain geographically limited to the upper reaches of the Oyodo River system and a tributary of the Tone River system (Ministry of the Environment, 2025; Tsuji et al., 2024). Therefore, most river sections identified in this study as having high habitat suitability potential are located in areas that are either still uninvaded or at an early stage of invasion.

Because management interventions are generally more cost-effective and more likely to succeed before populations become widely established (Simberloff et al., 2013), identifying such areas in advance may be particularly valuable. Although nationwide uniform monitoring across all river systems would be logistically challenging given the spatial extent of Japan’s freshwater networks, risk-based prioritisation informed by SDM outputs should support more strategic allocation of limited resources (McGeoch et al., 2016b).

### Strategies to prevent spread through new and secondary releases

The finding that the Korean perch exhibits high habitat suitability potential across large parts of the Japanese archipelago highlights the need for maximum vigilance against further range expansion through new or secondary releases. In freshwater fishes such as the Korean perch, artificial translocation is widely recognised as one of the primary drivers of rapid range expansion, often enabling non-native species to overcome dispersal barriers that would otherwise limit natural dispersal (Rahel, 2007).

Because the Korean perch is popular both as an ornamental species and as a target for angling, there is a persistent risk of transport and release by individuals. Indeed, within the Oyodo River system, range expansion beyond weirs and dams several metres in height without fishways has been found at multiple sites, strongly suggesting the occurrence of artificial secondary releases (Tsuji et al., 2024; Tsuji et al. in prep.). Furthermore, in 2024, one individual of the Korean perch was detected in a tributary of the Tone River system located several hundred kilometres away from the Oyodo River basin, providing clear evidence of a new or secondary introduction event (Ministry of the Environment, 2025). In geographically distant regions from the Oyodo River system, natural dispersal is highly unlikely even where habitat suitability is high; however, human-mediated introductions could readily lead to invasion and establishment.

One of the most critical factors in preventing nationwide spread is preventing human-mediated movement of the species. Currently, the Korean perch is legally designated as a “designated invasive alien species” by the Ministry of the Environment, and the transport, keeping, and release of live individuals are prohibited. However, experiences with other predatory invasive fishes subject to comparable legal regulation, such as largemouth bass and bluegill, indicate that legal restrictions alone may not fully prevent continued spread when illegal releases occur (Hosoya, 2019; Iguchi et al., 2004; Maezono & Miyashita, 2003). While the social and ecological contexts of these species differ, these cases suggest that complementary measures beyond regulation may be necessary. In addition to legal regulation, strengthening outreach and educational activities targeted at residents, citizen scientists, anglers, and others is likely to be essential for preventing further spread of the Korean perch (Rahel, 2007). By clearly communicating the ecological impacts of invasive species, the risks associated with release, and the seriousness of such actions as illegal behaviour, it may be possible to reduce releases. Furthermore, providing information based on scientific findings, such as habitat suitability predictions, could enhance public awareness and vigilance regarding Korean perch, potentially increasing individual-level monitoring and early detection across river systems. Ultimately, the practical impact of these strategies would depend on their integration with field-based monitoring, stakeholder engagement, and adaptive management frameworks.

### Discrepancies between model predictions and observed establishment: why has establishment been successful?

Invasion studies have frequently reported that species may occupy environmental conditions in introduced regions that differ from those observed in their native range, a phenomenon described as niche shift or expansion (Broennimann et al., 2007; Guisan et al., 2014; Tingley et al., 2014). Under such circumstances, SDMs calibrated solely with native-range occurrence data may underestimate realised distribution or subsequent spread in the introduced region (Pili et al., 2020). Therefore, it is important to interpret the habitat suitability predicted in this study as reflecting environmental associations observed within the native range, rather than as a strict upper bound of invasion risk.

In the Oyodo River system, the Korean perch has established and expanded despite predicted suitability values that were generally low to moderate (0.2–0.6) (Hibino et al., 2022; Tsuji et al., 2024). First, correlative SDMs rely on the environmental variables included in the model; unmeasured abiotic factors or spatial resolution constraints may influence local establishment success. Second, the realised niche in the introduced range may differ from that observed in the native range due to changes in biotic interactions or ecological release. Third, stochastic processes and propagule pressure may facilitate establishment even in areas predicted as only moderately suitable. Biotic interactions are one plausible factor influencing establishment success. In the native range in South Korea, large predatory fishes such as *Siniperca scherzeri* have been reported to function as top predators within riverine food webs (Chae et al., 2019). Although direct suppression of Korean perch populations by these species has not been quantitatively demonstrated, their presence may contribute to top-down regulation or competitive interactions that constrain population density and distribution. In contrast, native top predators are limited in the Oyodo River system and other Japanese freshwater areas, with almost no predators inhabiting the same mid- and upper river zone as the Korean perch (Iguchi et al., 2004). The absence of such biotic interactions is likely to reduce predation and competitive pressures, thereby increasing population density and establishment success and enabling range expansion beyond the environmental limits assumed by the SDM. Additionally, it has been suggested that native prey species may exhibit limited behavioural or evolutionary adaptation to novel predators, potentially increasing the predation success of introduced species (Sih et al., 2010). Whether such mechanisms operate in the case of the Korean perch in Japan remains to be empirically tested, but they represent a plausible hypothesis warranting further investigation.

While the SDM predicted only moderate habitat suitability for the tributary of the Tone River system (Ayu River), where a single individual was detected in 2024, follow-up surveys in 2025 recorded 101 individuals at the same river section, indicating successful establishment and rapid expansion (Ministry of the Environment, 2025; https://kanto.env.go.jp/topics_00414.html). This newly available evidence indicates that even areas predicted to have moderate suitability may support substantial population growth once introduction has occurred. Habitat suitability was predicted to increase towards upstream sections of this tributary, suggesting that environmental conditions may become progressively more favourable along the longitudinal gradient. Should upstream habitats combine higher suitability with continued propagule pressure or downstream dispersal, further range expansion within the Tone River system may occur, as in the Oyodo River system. The observed rapid increase in abundance within a short time frame underscores the importance of timely intervention. Invasion appears to be in an early but accelerating phase in this river system. Under such conditions, early and intensive eradication or containment measures, implemented before widespread dispersal occurs, may offer a greater likelihood of success compared with delayed responses.

### Strengths, limitations, and future directions of SDM-based invasion risk assessment

The RF–based SDM developed in this study demonstrated acceptable predictive performance, with AUC values of 0.86 for the training dataset and 0.80 for the independent test dataset, and a TSS value of 0.56. These values indicate reasonable discrimination ability between suitable and unsuitable conditions. In addition, predicted high-suitability areas broadly corresponded to regions with dense occurrence records in the native range, supporting the ecological plausibility of the model outputs. Spatial cross-validation further indicated that model performance remained stable across environmentally distinct catchment-level partitions, suggesting a degree of generalisability. Nevertheless, as with all machine learning approaches, random forest models are susceptible to overfitting if model complexity is not appropriately constrained (Elith et al., 2008; Olden et al., 2008). Particularly for random forest models, their high flexibility means that default settings tend to produce overly complex decision boundaries (Breiman, 2001). In the present study, steps including down-sampling, adjustment of node-splitting criteria, constraints on tree depth, and ensemble averaging were implemented to reduce this risk. The relatively small difference between training and test performance suggests that severe overfitting was unlikely, although it cannot be entirely excluded.

In the context of the Korean perch invasion examined here, an important strength of SDMs lies in their ability to provide spatially explicit estimates of environmental suitability across broad geographic extents (Thuiller et al., 2005). Conducting detailed surveys and management across all regions is impractical; therefore, decisions on where and to what extent to implement measures are constantly required within limited resources. SDMs provide valuable scientific and practical support for field-level decision-making. However, as discussed above, it is essential to recognise that correlative SDMs primarily quantify relationships between species occurrences and abiotic environmental variables (Araújo & Guisan, 2006; Elith & Leathwick, 2009). They do not explicitly incorporate propagule pressure, biotic interactions (e.g. predation and competition), dispersal dynamics, demographic processes, or temporal lags in establishment. Future advances in SDM-based invasion risk assessment would therefore benefit from integrative approaches that combine environmental suitability modelling with data on introduction pathways, population dynamics during early invasion stages, and biotic interaction networks (Yates et al., 2018; Zurell et al., 2016).

## Acknowledgements

We thank Dr. Hideyuki Doi, Dr. Katsutoshi Watanabe and Dr. Kei Hiroi (Kyoto university) for their accurate and valuable advice on this study. All experiments in the present study were compliant with the current laws of the country, Japan, in which they were performed.

## Statements and Declarations

We declare no conflicts of interest.

## Competing interests

Not applicable

## Funding

This study was supported by JSPS KAKENHI Grant Number 23K13967.

## Authors’ contributions

**S.M.**; Methodology, Modelling, Writing − review & editing: **S.T**.; Conceptualisation, Methodology, Writing − original draft & editing, Funding acquisition, Supervision

## Figure captions

Fig. S1 Correlation matrix between environmental variables. Each value indicates Pearson’s correlation coefficient (r), with colour denoting the strength and sign of the correlation

Fig. S2 (a) Basin boundaries (grey lines) and major rivers (blue lines) based on HydroBASINS Level 8 in South Korea. (b) Schematic diagram of nested spatial cross-validation using basin boundaries from HydroBASINS level 8. In the outer split, the training dataset (blue/orange) and the validation dataset (red) were partitioned at the basin level. In the inner split, five-fold cross-validation was performed at the basin level using the training dataset. Orange indicates the validation data for each iteration

## Notes

### Competing Interest Statement

The authors have declared no competing interest.

## References

Araújo, M. B., & Guisan, A. (2006). Five (or so) challenges for species distribution modelling. Journal of Biogeography, 33(10), 1677–1688. 10.1111/j.1365-2699.2006.01584.x

Breiman, L. (2001). Random Forests. Machine Learning, 45(1), 5–32. 10.1023/A:1010933404324

Broennimann, O., Treier, U. A., Müller-Schärer, H., Thuiller, W., Peterson, A. T., & Guisan, A. (2007). Evidence of climatic niche shift during biological invasion. Ecology Letters, 10(8), 701–709. 10.1111/j.1461-0248.2007.01060.x

Byeon, H.-K. (2017). Studies on the Feeding Habits of Korean aucha perch, *Coreoperca herzi* in the Geum River, Korea. Korean Journal of Environment and Ecology, 31, 472–478. 10.13047/KJEE.2017.31.5.472

Chae, B., Song, H., & Park, J. (2019). A field guide to the freshwater fishes of Korea. LG Evergreen Foundation.

Church, M. (2002). Geomorphic thresholds in riverine landscapes. Freshwater Biology, 47(4), 541–557. 10.1046/j.1365-2427.2002.00919.x

Cox, J. G., & Lima, S. L. (2006). Naiveté and an aquatic–terrestrial dichotomy in the effects of introduced predators. Trends in Ecology & Evolution, 21(12), 674–680. 10.1016/j.tree.2006.07.011

Cutler, D. R., Edwards Jr., T. C., Beard, K. H., Cutler, A., Hess, K. T., Gibson, J., & Lawler, J. J. (2007). Random Forests for Classification in Ecology. Ecology, 88(11), 2783–2792. 10.1890/07-0539.1

Diagne, C., Leroy, B., Vaissière, A.-C., Gozlan, R. E., Roiz, D., Jarić, I., Salles, J.-M., Bradshaw, C. J. A., & Courchamp, F. (2021). High and rising economic costs of biological invasions worldwide. Nature, 592(7855), 571–576. 10.1038/s41586-021-03405-6

Dukes, J. S., & Mooney, H. A. (1999). Does global change increase the success of biological invaders? Trends in Ecology & Evolution, 14(4), 135–139. 10.1016/S0169-5347(98)01554-7

Early, R., Bradley, B. A., Dukes, J. S., Lawler, J. J., Olden, J. D., Blumenthal, D. M., Gonzalez, P., Grosholz, E. D., Ibañez, I., Miller, L. P., Sorte, C. J. B., & Tatem, A. J. (2016). Global threats from invasive alien species in the twenty-first century and national response capacities. Nature Communications, 7(1), 12485. 10.1038/ncomms12485

Elith, J., H. Graham*, C., P. Anderson, R., Dudík, M., Ferrier, S., Guisan, A., J. Hijmans, R., Huettmann, F., R. Leathwick, J., Lehmann, A., Li, J., G. Lohmann, L., A. Loiselle, B., Manion, G., Moritz, C., Nakamura, M., Nakazawa, Y., McC. M. Overton, J., Townsend Peterson, A., … E. Zimmermann, N. (2006). Novel methods improve prediction of species’ distributions from occurrence data. Ecography, 29(2), 129–151. 10.1111/j.2006.0906-7590.04596.x

Elith, J., Kearney, M., & Phillips, S. (2010a). The art of modelling range-shifting species. Methods in Ecology and Evolution, 1(4), 330–342. 10.1111/j.2041-210X.2010.00036.x

Elith, J., Kearney, M., & Phillips, S. (2010b). The art of modelling range-shifting species. Methods in Ecology and Evolution, 1(4), 330–342. 10.1111/j.2041-210X.2010.00036.x

Elith, J., & Leathwick, J. R. (2009). Species Distribution Models: Ecological Explanation and Prediction Across Space and Time. Annual Review of Ecology, Evolution, and Systematics, 40(Volume 40, 2009), 677–697. 10.1146/annurev.ecolsys.110308.120159

Elith, J., Leathwick, J. R., & Hastie, T. (2008). A working guide to boosted regression trees. Journal of Animal Ecology, 77(4), 802–813. 10.1111/j.1365-2656.2008.01390.x

Faghihinia, M., Xu, Y., Liu, D., & Wu, N. (2021). Freshwater biodiversity at different habitats: Research hotspots with persistent and emerging themes. Ecological Indicators, 129, 107926. 10.1016/j.ecolind.2021.107926

Fick, S. E., & Hijmans, R. J. (2017). WorldClim 2: New 1-km spatial resolution climate surfaces for global land areas. International Journal of Climatology, 37(12), 4302–4315. 10.1002/joc.5086

Gallardo, B., Clavero, M., Sánchez, M. I., & Vilà, M. (2016). Global ecological impacts of invasive species in aquatic ecosystems. Global Change Biology, 22(1), 151–163. 10.1111/gcb.13004

Gippet, J. M., Liebhold, A. M., Fenn-Moltu, G., & Bertelsmeier, C. (2019). Human-mediated dispersal in insects. *Current Opinion in Insect Science*, Global Change Biology • Molecular Physiology, 35, 96–102. 10.1016/j.cois.2019.07.005

Guisan, A., Petitpierre, B., Broennimann, O., Daehler, C., & Kueffer, C. (2014). Unifying niche shift studies: Insights from biological invasions. Trends in Ecology & Evolution, 29(5), 260–269. 10.1016/j.tree.2014.02.009

Guisan, A., & Thuiller, W. (2005). Predicting species distribution: Offering more than simple habitat models. Ecology Letters, 8, 909–1020. 10.1111/j.1461-0248.2005.00792.x

Guisan, A., & Zimmermann, N. E. (2000). Predictive habitat distribution models in ecology. Ecological Modelling, 135(2), 147–186. 10.1016/S0304-3800(00)00354-9

Havel, J. E., Kovalenko, K. E., Thomaz, S. M., Amalfitano, S., & Kats, L. B. (2015). Aquatic invasive species: Challenges for the future. Hydrobiologia, 750(1), 147–170. 10.1007/s10750-014-2166-0

Hibino, Y., Ogata, Y., Matsuo, R., Ooe, R., Obaru, N., Kurihara, T., & Saiki, Y. (2022). Expansion of Korean perch *Coreoperca herzi* in the Oyodo River system, and presumed impact of the fish on native fishes. *Ichthy*, Natural History of Fishes of Japan, 16, 18–24.

Hibino, Y., Taguchi, T., Iwata, K., & Furuhashi, R. (2019). Records of *Coreoperca herzi* (Perciformes: Sinipercidae) from Oyodo River system, Miyazaki Prefecture, Japan. Nature of Kagoshima, 45, 243–248.

Hosoya, K. (2019). Yamakei Handy Illustrated Book 15: Freshwater fishes of Japan. Yama-Kei Publishers.

Iguchi, K., Matsuura, K., McNyset, K. M., Peterson, A. T., Scachetti-Pereira, R., Powers, K. A., Vieglais, D. A., Wiley, E. O., & Yodo, T. (2004). Predicting Invasions of North American Basses in Japan Using Native Range Data and a Genetic Algorithm. Transactions of the American Fisheries Society, 133(4), 845–854. 10.1577/T03-172.1

Lehner, B., & Grill, G. (2013). Global river hydrography and network routing: Baseline data and new approaches to study the world’s large river systems. Hydrological Processes, 27(15), 2171–2186. 10.1002/hyp.9740

Leonardi, M., Colucci, M., Pozzi, A. V., Scerri, E. M. L., & Manica, A. (2024). tidysdm: Leveraging the flexibility of tidymodels for species distribution modelling in R. Methods in Ecology and Evolution, 15(10), 1789–1795. 10.1111/2041-210X.14406

Linke, S., Lehner, B., Ouellet Dallaire, C., Ariwi, J., Grill, G., Anand, M., Beames, P., Burchard-Levine, V., Maxwell, S., Moidu, H., Tan, F., & Thieme, M. (2019). Global hydro-environmental sub-basin and river reach characteristics at high spatial resolution. Scientific Data, 6(1), 283. 10.1038/s41597-019-0300-6

Maezono, Y., & Miyashita, T. (2003). Community-level impacts induced by introduced largemouth bass and bluegill in farm ponds in Japan. Biological Conservation, 109(1), 111–121. 10.1016/S0006-3207(02)00144-1

McEachran, M. C., Hofelich Mohr, A., Lindsay, T., Fulton, D. C., & Phelps, N. B. D. (2022). Patterns of live baitfish use and release among recreational anglers in a regulated landscape. North American Journal of Fisheries Management, 42(2), 295–306. 10.1002/nafm.10747

McGeoch, M. A., Genovesi, P., Bellingham, P. J., Costello, M. J., McGrannachan, C., & Sheppard, A. (2016a). Prioritizing species, pathways, and sites to achieve conservation targets for biological invasion. Biological Invasions, 18(2), 299–314. 10.1007/s10530-015-1013-1

McGeoch, M. A., Genovesi, P., Bellingham, P. J., Costello, M. J., McGrannachan, C., & Sheppard, A. (2016b). Prioritizing species, pathways, and sites to achieve conservation targets for biological invasion. Biological Invasions, 18(2), 299–314. 10.1007/s10530-015-1013-1

Ministry of the Environment. (2025). Confirmation of the non-native species Korean perch in the Tone River System. https://ikilog.biodic.go.jp/files/eDNA_Coreoperca.pdf

Olden, J. D., Lawler, J. J., & Poff, N. L. (2008). Machine Learning Methods Without Tears: A Primer for Ecologists. The Quarterly Review of Biology, 83(2), 171–193. 10.1086/587826

Pili, A. N., Tingley, R., Sy, E. Y., Diesmos, M. L. L., & Diesmos, A. C. (2020). Niche shifts and environmental non-equilibrium undermine the usefulness of ecological niche models for invasion risk assessments. Scientific Reports, 10(1), 7972. 10.1038/s41598-020-64568-2

Poole, G. C., & Berman, C. H. (2001). An ecological perspective on in-stream temperature: Natural heat dynamics and mechanisms of human-caused thermal degradation. Environmental Management, 27(6), 787–802. 10.1007/s002670010188

Punzet, M., Voß, F., Voß, A., Kynast, E., & Bärlund, I. (2012). A global approach to assess the potential impact of climate change on stream water temperatures and related in-stream first-order decay rates. Journal of Hydrometeorology, 13(3), 1052–1065. 10.1175/JHM-D-11-0138.1

R Core Team. (2021). R: A Language and Environment for Statistical Computing.

Rahel, F. J. (2007). Biogeographic barriers, connectivity and homogenization of freshwater faunas: It’s a small world after all. Freshwater Biology, 52(4), 696–710. 10.1111/j.1365-2427.2006.01708.x

Rodríguez-Castañeda, G., Hof, A. R., Jansson, R., & Harding, L. E. (2012). Predicting the fate of biodiversity using species’ distribution models: Enhancing model comparability and repeatability. PLOS ONE, 7(9), e44402. 10.1371/journal.pone.0044402

Sato, M., Kawaguchi, Y., Nakajima, J., Mukai, T., Shimatani, Y., & Onikura, N. (2010). A review of the research on introduced freshwater fishes: New perspectives, the need for research, and management implications. Landscape and Ecological Engineering, 6(1), 99–108. 10.1007/s11355-009-0086-3

Seebens, H., Bacher, S., Blackburn, T. M., Capinha, C., Dawson, W., Dullinger, S., Genovesi, P., Hulme, P. E., van Kleunen, M., Kühn, I., Jeschke, J. M., Lenzner, B., Liebhold, A. M., Pattison, Z., Pergl, J., Pyšek, P., Winter, M., & Essl, F. (2021). Projecting the continental accumulation of alien species through to 2050. Global Change Biology, 27(5), 970–982. 10.1111/gcb.15333

Sih, A., Bolnick, D. I., Luttbeg, B., Orrock, J. L., Peacor, S. D., Pintor, L. M., Preisser, E., Rehage, J. S., & Vonesh, J. R. (2010). Predator–prey naïveté, antipredator behavior, and the ecology of predator invasions. Oikos, 119(4), 610–621. 10.1111/j.1600-0706.2009.18039.x

Simberloff, D., Martin, J.-L., Genovesi, P., Maris, V., Wardle, D. A., Aronson, J., Courchamp, F., Galil, B., García-Berthou, E., Pascal, M., Pyšek, P., Sousa, R., Tabacchi, E., & Vilà, M. (2013). Impacts of biological invasions: What’s what and the way forward. Trends in Ecology & Evolution, 28(1), 58–66. 10.1016/j.tree.2012.07.013

Thuiller, W., Richardson, D. M., Pyšek, P., Midgley, G. F., Hughes, G. O., & Rouget, M. (2005). Niche-based modelling as a tool for predicting the risk of alien plant invasions at a global scale. Global Change Biology, 11(12), 2234–2250. 10.1111/j.1365-2486.2005.001018.x

Tingley, R., Vallinoto, M., Sequeira, F., & Kearney, M. R. (2014). Realized niche shift during a global biological invasion. Proceedings of the National Academy of Sciences, 111(28), 10233–10238. 10.1073/pnas.1405766111

Tsuji, S., Doi, H., Hibino, Y., Shibata, N., & Watanabe, K. (2024). Rapid assessment of invasion front and biological impact of the invasive fish *Coreoperca herzi* using quantitative eDNA metabarcoding. Biological Invasions, 26(9), 3107–3123. 10.1007/s10530-024-03364-9

Turbelin, A. J., Hudgins, E. J., Catford, J. A., Cuthbert, R. N., Diagne, C., Kourantidou, M., Roiz, D., & Courchamp, F. (2024). Biological invasions as burdens to primary economic sectors. Global Environmental Change, 87, 102858. 10.1016/j.gloenvcha.2024.102858

Uchida, K. (1935). Life history of *Coreoperca herzi*. Zoological Magazine, 47, 257–275.

Valavi, R., Elith, J., Lahoz-Monfort, J. J., & Guillera-Arroita, G. (2021). Modelling species presence-only data with random forests. Ecography, 44(12), 1731–1742. 10.1111/ecog.05615

Valavi, R., Elith, J., Lahoz-Monfort, J. J., & Guillera-Arroita, G. (2023). Flexible species distribution modelling methods perform well on spatially separated testing data. Global Ecology and Biogeography, 32(3), 369–383. 10.1111/geb.13639

Valavi, R., Guillera-Arroita, G., Lahoz-Monfort, J. J., & Elith, J. (2022). Predictive performance of presence-only species distribution models: A benchmark study with reproducible code. Ecological Monographs, 92(1), e01486. 10.1002/ecm.1486

Wright, M. N., & Ziegler, A. (2017). ranger: A fast implementation of random forests for high dimensional data in C++ and R. Journal of Statistical Software, 77, 1–17. 10.18637/jss.v077.i01

Yates, K. L., Bouchet, P. J., Caley, M. J., Mengersen, K., Randin, C. F., Parnell, S., Fielding, A. H., Bamford, A. J., Ban, S., Barbosa, A. M., Dormann, C. F., Elith, J., Embling, C. B., Ervin, G. N., Fisher, R., Gould, S., Graf, R. F., Gregr, E. J., Halpin, P. N., … Sequeira, A. M. M. (2018). Outstanding challenges in the transferability of ecological models. Trends in Ecology & Evolution, 33(10), 790–802. 10.1016/j.tree.2018.08.001

Zurell, D., Thuiller, W., Pagel, J., Cabral, J. S., Münkemüller, T., Gravel, D., Dullinger, S., Normand, S., Schiffers, K. H., Moore, K. A., & Zimmermann, N. E. (2016). Benchmarking novel approaches for modelling species range dynamics. Global Change Biology, 22(8), 2651–2664. 10.1111/gcb.13251

