## Supplementary figures and images for "Predicting establishment risk of the Korean perch *Coreoperca herzi* in Japan using species distribution modelling: Insights for invasive fish management"

### Supplemental figures

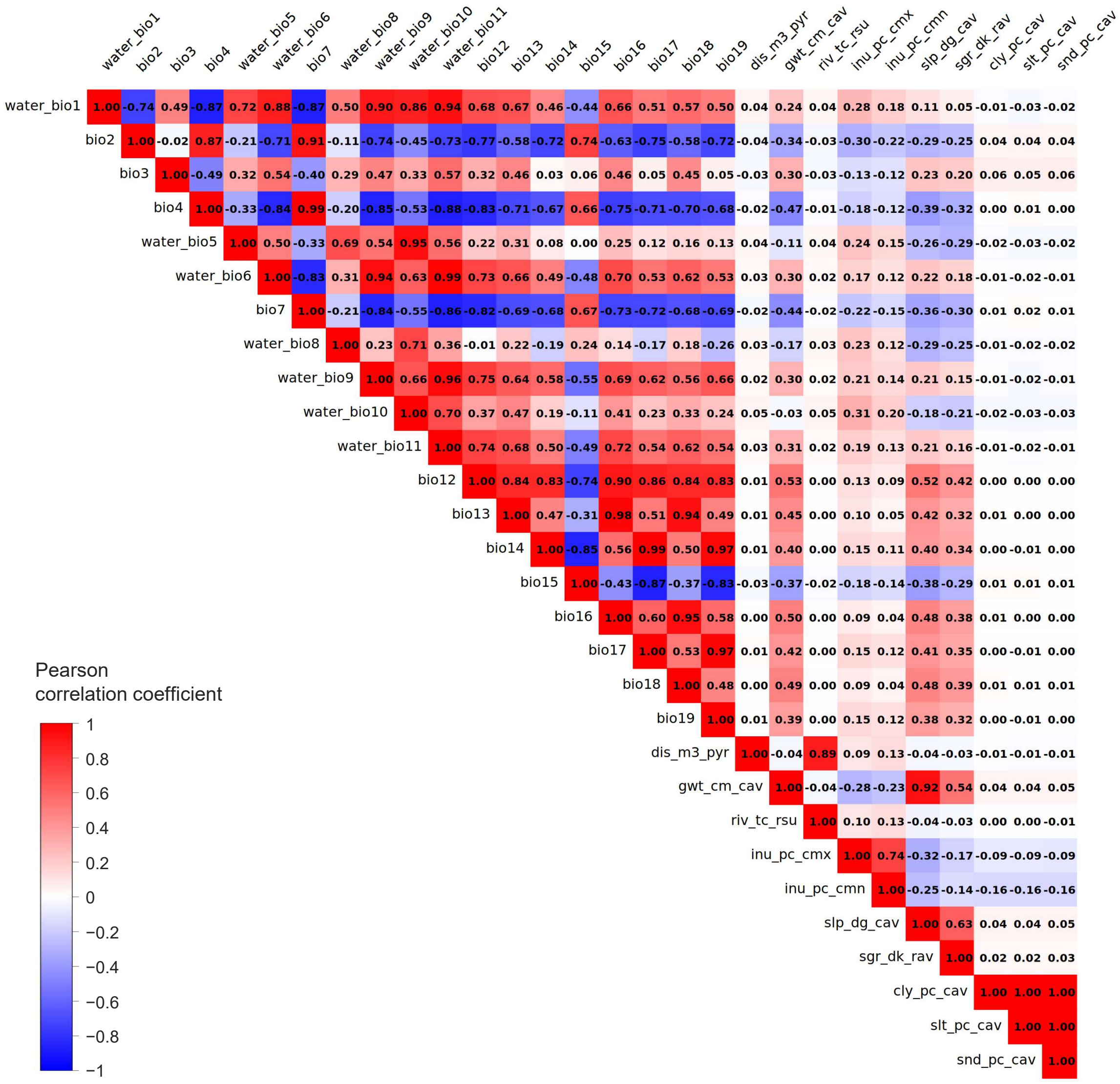

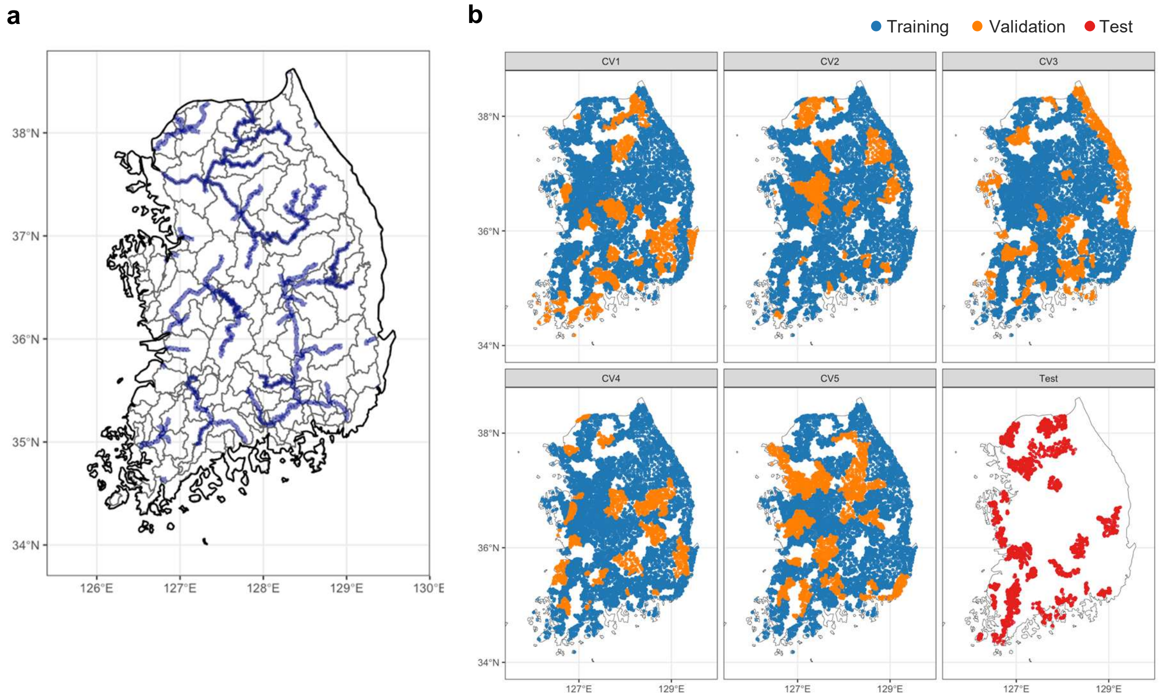
